# Blebbistatin protects iPSC-CMs from hypercontraction and facilitates automated patch-clamp based electrophysiological study

**DOI:** 10.1101/2021.06.07.447362

**Authors:** Wener Li, Xiaojing Luo, Ying Ulbricht, Kaomei Guan

## Abstract

Recently, there have been great advances in cardiovascular channelopathy modeling and drug safety pharmacology using human induced pluripotent stem cell-derived cardiomyocytes (iPSC-CMs). The automated patch-clamp (APC) technique overcomes the disadvantages of the manual patch-clamp (MPC) technique, which is labor intensive and gives low output. However, the application of the APC platform is still limited in iPSC-CM based research, due to the difficulty in maintaining the high quality of single iPSC-CMs during dissociation and recording. In this study, we improved the method for single iPSC-CM preparation by applying 2.5 μM blebbistatin (BB, an excitation-contraction coupling uncoupler) throughout APC procedures (dissociation, filtration, storage, and recording). Under non-BB buffered condition, iPSC-CMs in suspension showed a severe bleb-like morphology. However, BB-supplement led to significant improvements in morphology and *I*_*Na*_ recording, and we even obtained several CMs that showed spontaneous action potentials with typical morphology. Furthermore, APC faithfully recapitulated the single-cell electrophysiological phenotypes of iPSC-CMs derived from Brugada syndrome patients, as detected with MPC. Our study indicates that APC is capable of replacing MPC in the modeling of cardiac channelopathies using human iPSC-CMs by providing high-quality data with higher throughput.

## Introduction

Since the development of patch-clamp techniques for high-resolution current recording from cells and cell-free membrane patches, the manual patch-clamp (MPC) technology revolutionized electrophysiological studies [1]. The high quality of the data obtained with the MPC technique is highly appreciated, but the throughput with this technique is notoriously low [2]. So far, the modelling of cardiac channelopathies using human induced pluripotent stem cell-derived cardiomyocytes (iPSC-CMs) is mostly achieved by studying ion channels using MPC. For example, MPC has been widely used in recordings of *I*_*Na*_ (voltage-gated sodium current), *I*_*CaL*_ (L-type voltage-gated calcium current), or *I*_*kr*_ (the rapid component of the delayed rectifier potassium current) in iPSC-CMs to study cardiac channelopathies caused by mutations in genes encoding ion channel proteins, such as *SCN5A* [3], *CACNA1C* [4] or *KCNH2* [5, 6]. For a better overview of the use of MPC in iPSC-CM models of channelopathies, please see the recently published review [7]. However, the labor-intensive and low-throughput nature of MPC has hindered its large implementation in drug discovery. With these advantages and limitations in mind, the automated patch-clamp (APC) technique might play a key role in ion channel research as well as in drug discovery and safety testing [8]. However, APC poses an often underestimated challenge to the reproducibility of the data with iPSC-CMs, which are obtained using MPC [2].

The most challenging step for APC is the acquisition of a large number of single iPSC-CMs in suspension at the relaxation stage [9]. It is assumed that the Ca^2+^-free period is necessary to allow cell separation via the disruption of the Ca^2+^-dependent cadherins [10]. However, after only a few minutes of Ca^2+^-free perfusion and Ca^2+^ repletion, the profound changes to the cardiomyocytes, including ultrastructural alterations, loss of intracellular components, and Na^+^ and Ca^2+^ gain, were reported [11]. This phenomenon was first reported as the calcium paradox more than 50 years ago [12]. Since then, many studies have focused on a better understanding of Ca^2+^-paradox injury [11, 13-15]. To conquer this paradox, excitation-contraction uncoupling agents like 2,3-butanedione monoxime (BDM) or blebbistatin (BB) were commonly used for the isolation of cardiomyocytes from the Langendorff perfused heart or heart tissue using the chunk-method [10]. Nevertheless, the BDM was reported to have side effects such as reducing *I*_*to*_ [16], attenuating β-adrenergic response of *I*_*CaL*_ [17], and even inhibiting mitochondrial respiration [18]. On the other hand, BB was rarely reported to have electrophysiological side effects in rodents [19, 20], but significant effects in the isolated rabbit heart [21].

In this study, we improved our previously published cell dissociation method suitable for APC [9] by supplementing 2.5 μM BB in the papain-EDTA-based dissociation solution. Additionally, 2.5 μM BB was also used for strainer filtration and storage of iPSC-CMs. The extracellular solutions in the liquid handling part of the APC system were also supplemented with 2.5 μM BB. We assume that the protection of fragile iPSC-CMs in suspension by applying BB would be beneficial for electrophysiological studies of ion channels by using APC. Our data demonstrate that BB (2.5 μM) itself does not exhibit any side effects on action potential (AP) and field potential (FP) using human iPSC-CMs. Importantly, BB protects human iPSC-CMs in suspension from Ca^2+^-paradox injury and enables recordings of ionic currents and spontaneous APs with APC. Furthermore, we conducted ion channel studies in iPSC-CMs from patients with Brugada syndrome (BrS) by using APC. The data on *I*_*Na*_, *I*_*to*_ and *I*_*CaL*_ were consistent with our previous data obtained by using MPC. Our study indicates that APC is capable of replacing MPC in the modeling of cardiac channelopathies using human iPSC-CMs by providing high quality data with higher throughput.

## Material and methods

### Directed differentiation of iPSCs into iPSC-CMs

Directed differentiation of iPSCs into ventricular-like CMs was induced by modulating WNT signaling as previously described [22]. In this study, we used three different human iPSC lines (iWT.D2-1/3, iBM76-1/3 and isWT1.Bld2), which were described previously [22]. When iPSCs (cultured on 12-well plates) reached around 90% confluency, differentiation was initiated by changing medium into cardio differentiation medium (RPMI 1640 with GlutaMax and HEPES (Thermo Fisher Scientific), 0.5 mg/ml human recombinant albumin (Sigma-Aldrich) and 0.2 mg/ml L-ascorbic acid 2-phosphate (Sigma-Aldrich)) supplemented with 4 μM of the GSK3β inhibitor CHIR99021 (Millipore). After 48 hours, the medium was changed to fresh cardio differentiation medium supplemented with 5 μM IWP2 (WNT signaling inhibitor, Millipore) for another two days. Afterwards, cells were cultured in the cardio differentiation medium for another 4 days. From day eight on, the cardio differentiation medium was replaced by RPMI/B27 medium (RPMI 1640 with GlutaMax and HEPES, supplemented with 2% B27 with insulin (Thermo Fisher Scientific)). On day 20, beating cardiomyocytes were detached from plates with 1 mg/ml collagenase B (Worthington Biochemical), dissociated with 0.25% Trypsin/EDTA (Thermo Fisher Scientific), and replated into Geltrex-coated 6-well plates at a density of 800,000 cells/well. Afterwards, iPSC-CMs were cultured in RPMI/B27 medium until 3 months.

### Dissociation of 3-month-old iPSC-CMs into single cells for automated patch-clamp

Our previously published dissociation method was used in this study with some modifications [9]. Collagenase B (1 mg/ml) was used to pre-treat 3-month-old iPSC-CMs until the layer of cardiomyocytes detached from the plate. The layer of cardiomyocytes was transferred into a 3.5-cm dish and then treated with 2 ml of 20 U/ml papain (Sigma-Aldrich) dissolved in 1.1 mM EDTA-buffered RPMI/B27 medium containing 2.5 μM BB (Sigma-Aldrich, dissolved in DMSO as a 10 mM stock) for 10 min. A fire-polished glass Pasteur pipette was used to gently agitate the cells to release single iPSC-CMs. The cell suspension was filtered through a 30-μm strainer (MACS SmartStrainers, Miltenyi Biotec) to remove cell clusters and was centrifuged for 1 min at 50 *g*. After gently aspirating the supernatant, the cell pellet was resuspended into 2 ml of 1.1 mM EDTA-buffered RPMI/B27 medium with 2.5 μM BB and then filtered through a 10-μm strainer (pluriStrainer^®^, pluriSelect) to collect larger iPSC-CMs. The cells were collected with 2 ml of RPMI/B27 medium with 2.5 μM BB and then centrifuged for 1 min at 50 *g*. After removing the supernatant, the cell pellet was gently resuspended and further stored in 2.5 μM BB-containing RPMI/B27 medium at 4 °C for 2 h.

### Contractility and FP measurements

For the measurements of contractility and FP together in the same culture, the Maestro Edge multiwell microelectrode array (MEA) and impedance system (Axion BioSystems) was used. The 3-month-old iPSC-CMs were dissociated with 0.25% Trypsin-EDTA and seeded at a density of 10,000 cells in 8 μl of digestion medium (RPMI/B27 with 20% fetal calf serum, and 1 μM Thiazovivin) into Cytoview MEA 6-well (with 64 electrodes) or 24-well (with 16 electrodes) plates for 1 hour (Axion BioSystems). After cells attached to the plate, 1 ml of digestion medium was added to each well. Every well was pre-coated with 8 μl Geltrex^®^ (Thermo Fisher Scientific) for at least 1 hour before use. The medium was changed to RPMI/B27 medium one day after plating and thereafter every two days until day 6. On day 6, electrical FP and the impedance-based contractility in human iPSC-CMs were measured before and after 2.5 μM BB treatment. After calibrating for 10 minutes, the spontaneous recordings were carried out at 37 °C and 5% CO_2_ using the AxIS Navigator software (Axion BioSystems). The sample rates were 12,500 Hz for FP and 40 Hz for contractility. Spontaneous beating frequency was defined by the reciprocal of averaged inter-beat interval. The spontaneous beating frequency, FP amplitude, and conduction velocity were generated by using the AxIS Navigator software and further analyzed using AxIS Metric Plotting Tool (Axion BioSystems). The mainstream conduction velocity values were averaged for one culture.

### Manual patch-clamp for AP measurement

Human iPSC-CMs around day 90 were enzymatically dissociated into single cells and seeded on 5 mm Ø coverslips distributed in 35-mm dishes. After around 10 days for recovery, the paced APs of a single iPSC-CM were measured at room temperature with a ruptured whole-cell current clamp using HEKA EPC10 amplifier and Patchmaster (HEKA Elektronik). The pipette and extracellular solutions for paced AP recordings were listed in Supplementary Table 1. The pacing stimulus was 0.5 Hz. Pipette potentials were corrected for liquid junction potentials. More than 5 continuously stable, paced APs were chosen and analyzed using LabChart^®^ (ADInstruments) to determine AP duration at 90% repolarization (APD90), AP amplitude (APA), and resting membrane potential (RMP).

### Calcium transient measurement

Paced whole-cell calcium transients were measured according to our previous publication [23]. CMs around day 80 were dissociated and replated on coverslips at a density of 200,000 cells/well (6-well plate). Cells recovered for at least 10 days were loaded with Fura-2 (Thermo Fisher Scientific) at a final concentration of 5 μM in RPMI/B27 medium for 30 min at 37 °C and washed twice with the medium. Before measurement, cells were incubated for 10 min to enable complete de-esterification of intracellular Fura-2. Calcium transients were recorded using a 40× objective on an Olympus IX70 microscope fitted with an IonWizard software (IonOptix) at 35 °C. Samples were excited at 340 and 380 nm with a switching frequency of 200 Hz and the emitted fluorescence was collected at 510 nm. The cytosolic Ca^2+^ level was measured as the ratio of fluorescence at 340 and 380 nm (340/380 nm) in Tyrode’s solution. To minimize the phototoxicity and photoinactivation effects of BB, the recording was paused during the BB exposure time. To normalize the Ca^2+^ transient frequency, iPSC-CMs were field-stimulated using a MyoPacer (IonOptix) at a pacing frequency of 0.25 Hz (6 V, 10 ms). The monotonic transient analysis was performed using LabChart^®^ (ADInstruments) and the following parameters were determined: peak amplitude of Ca^2+^ transients (the Fura-2 ratio at systole subtracted by the Fura-2 ratio at diastole), decay rate (tau), as well as duration of Ca^2+^ transients.

### Automated patch-clamp

All experiments were performed at room temperature using an APC system (Patchliner Quattro, Nanion Technologies GmbH) with low resistance NPC-16 chips. The pipette and extracellular solutions for *I*_*Na*_, *I*_*to*_, and *I*_*CaL*_ recordings were listed in Supplementary Table 1. From a holding potential of −100 mV, *I*_*Na*_ was recorded using voltage steps from −80 to +70 mV for 20 ms in a 5 mV step at an interval of 2000 ms (shown as an inset in Figure 3B and 4B). Nifedipine (10 μM) was used to block *I*_*CaL*_. *I*_*to*_ was recorded by increasing the testing potential stepwise from −40 mV to +60 mV in a 10 mV step from a holding potential of −90 mV with a 20 ms pre-pulse to −35 mV to inactivate *I*_*Na*_ (shown as an inset in Figure 4D). CdCl_2_ (0.5 mM) was used to block calcium current. Each pulse lasted for 400 ms, the sweep interval was 10 s. To record *I*_*CaL*_, cells were depolarized for 100 ms to voltages between −80 to +50 mV from a holding potential of −90 mV, the sweep interval was 3 s (shown as an inset in Figure 4F). Currents were sampled at 25 kHz and low-pass-filtered at 2.9 kHz. The cells with series resistance < 5 MΩ were used for further analysis. The liquid junction potentials were not compensated during recordings but adjusted after the measurement for each cell separately. The liquid junction potentials were assessed under current-clamp mode with the MPC system.

The dynamic patch-clamp experiment for AP recordings was carried out using the Dynamite system from Nanion GmbH [24]. *I*_*K1*_ conductance was set to 2000 pS/pF. The pacing pulse was 800 pA with a duration of 2 ms and at a rate of 0.5 Hz (shown as an inset in Figure 3E).

The data were exported by using Patchmaster and further analyzed with Graph Pad Prism 5 (GraphPad Software, Inc).

### Statistics

Statistical analysis was performed with GraphPad Prism 5 using the paired Student’s *t*-test to compare differences between two paired groups, and the two-way ANOVA with Bonferroni post-test for comparison of more groups and conditions. Data are presented as the mean ± standard error of the mean (SEM). For a better visualization of the spread of the data, boxplots were used in some experiments. Results were considered statistically significant when the p-value was < 0.05.

## Results and discussion

### Blebbistatin prevents freshly isolated iPSC-CMs from hypercontraction

In the previous study, we reported a Ca^2+^-free method for iPSC-CM dissociation into a single CM suspension [9]. We can obtain a large quantity of flake- and rod-like CMs and store the cells at the relaxation stage for at least 2 hours, which is required for patch-clamp experiments (Figure 1A). However, we observed the calcium paradox when we switch the solution from 1.1 mM EDTA-buffered RPMI/B27 medium to the physiological external solution (containing 2 mM CaCl_2_), which is used for APC. Almost all CMs immediately developed severe membrane blebs, no matter whether they were stored for 2 hours or only 5 minutes (Figure 1B).

**Figure 1.**
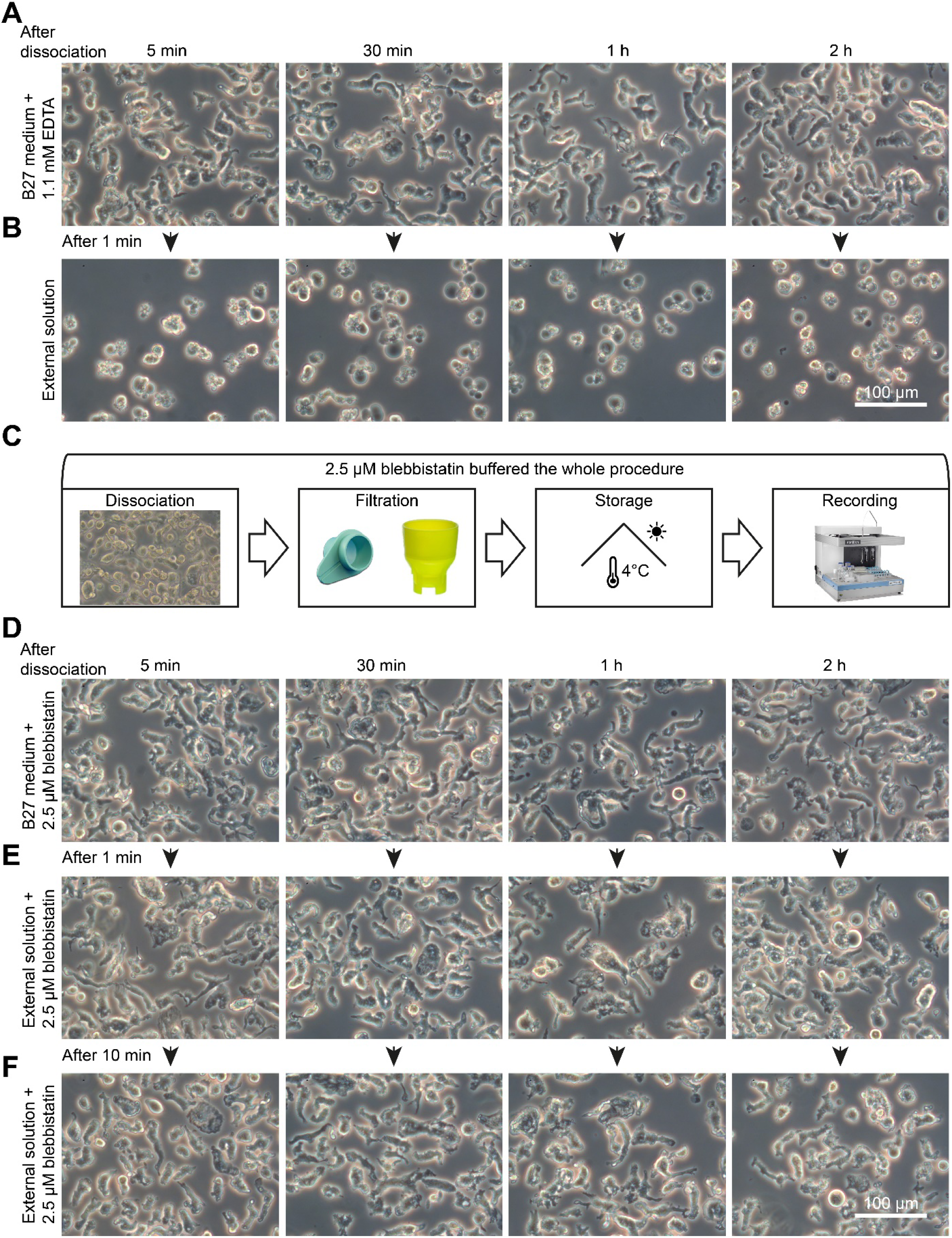
Morphological observations of dissociated iPSC-CMs in suspension under different conditions. (**A**) The morphologies of iPSC-CMs in suspension in 1.1 mM EDTA-buffered RPMI/B27 medium for 5 min, 30 min, 1 hour, and 2 hours. (**B**) The morphological changes after the CMs are transferred from EDTA-buffered RPMI/B27 medium to 2 mM Ca^2+^-containing physiological external solution, corresponding to the 4 different time points. (**C**) For the improved iPSC-CM preparation method, 2.5 μM BB was used to buffer the whole procedure including cell dissociation, filtration, storage, and recording. (**D**) The morphologies of iPSC-CMs in 2.5 μM BB-buffered RPMI/B27 medium for 5 min, 30 min, 1 hour, and 2 hours. (**E-F**) Correspondingly to different time points, shown are morphologies after the identical CMs transferred from BB-buffered RPMI/B27 medium to BB-buffered, 2 mM Ca^2+^-containing physiological external solution for 1 min (**E**) and 10 min (**F**). Scale bar: 100 μm.

The Ca^2+^-free period (1.1 mM EDTA-buffered RPMI/B27 medium) is necessary for CM separation by the disruption of the Ca^2+^-dependent cell-cell adhesion mediated by cadherins. However, previous study showed that switching from a Ca^2+^-free to a Ca^2+^-repletion condition used for CMs led to ultrastructural alterations, loss of intracellular components, and Na^+^ and Ca^2+^ gain [11]. To solve this calcium paradox issue, we tested the excitation-contraction uncoupler blebbistatin (BB, 2.5 μM, myosin II inhibitor [25]) in the whole APC procedure including the dissociation, filtration, storage, and recording steps (Figure 1C). Since BB is light sensitive [26], we avoid its light exposure during the whole procedure. We found the iPSC-CMs in suspension were maintained at the relaxation state even 2 hours after dissociation when the BB-buffered RPMI/B27 medium containing 0.4 mM Ca^2+^ was used as the storage solution (Figure 1D). To meet the needs of the APC process, where 2 mM Ca^2+^-containing physiological external solution was used for CM capturing, we switched from BB-buffered RPMI/B27 medium to BB-buffered external solution containing 2 mM CaCl_2_. We found that the majority of CMs remained the flake- and rod-like relaxation shapes for all 4 time points (5 min, 30 min, 1 h, and 2 hours) after approximately 1-minute of solution exchange (Figure 1E). To further test whether the flake- and rod-like shapes could persist throughout the patch progress, generally about 10 minutes, we recorded the CM morphologies and confirmed that the flake-and rod-like shapes were maintained for at least 10 minutes in BB-buffered physiological external solution (Figure 1F). Our data demonstrate that BB can be used for preparation of single cell suspension and storage of human iPSC-CMs. This is consistent with previous studies showing that BB can be used for the isolation and culture of high quality and viable adult mouse CMs [18, 27].

### Blebbistatin shows no effects on electrical signals of iPSC-CMs

Since the electrophysiological effects of BB have been controversially reported in different species, we assessed BB effects on contractility, FP, and AP in human iPSC-CMs. By utilizing the Maestro Edge multiwell MEA and impedance system, we could record the FP and impedance-based contractility in the same culture of iPSC-CMs before and after BB treatment for 10 min (Figure 2A-C). We tested 3 different concentrations of BB (1, 2.5 and 10 μM) and found that the contractility was abolished in most iPSC-CM cultures after the application of 2.5 μM BB for 3-5 minutes (Figure 2A, B), which is mainly due to the binding of BB to myosin II. However, the FP metrics such as spontaneous excitation frequency (Hz), spike amplitude (mV), and conduction velocity (cm/s) did not alter (Figure 2B and C). Similar results were obtained with 10 μM BB treatment, abolishing contraction within 1 minute (Supplementary Figure 1B). Treatment with 1 μM BB failed to completely block the contractility of iPSC-CMs within 10 minutes (Supplementary Figure 1A). Moreover, with the prolonged incubation of human iPSC-CMs with 2.5 μM BB up to 1 hour, we did not observe differences in FP parameters (Supplementary Figure 1C-E). These data indicate that the BB treatment (2.5 μM) has no effects on electrical field potential of human iPSC-CMs but prevents their beating. Our study is consistent with previous studies, in which the treatments with 1 μM [28] and 10 μM [29] BB did not show any effects on FP of human iPSC-CMs. Based on these findings, we applied 2.5 μM BB during the whole APC procedure for the following experiments.

**Figure 2.**
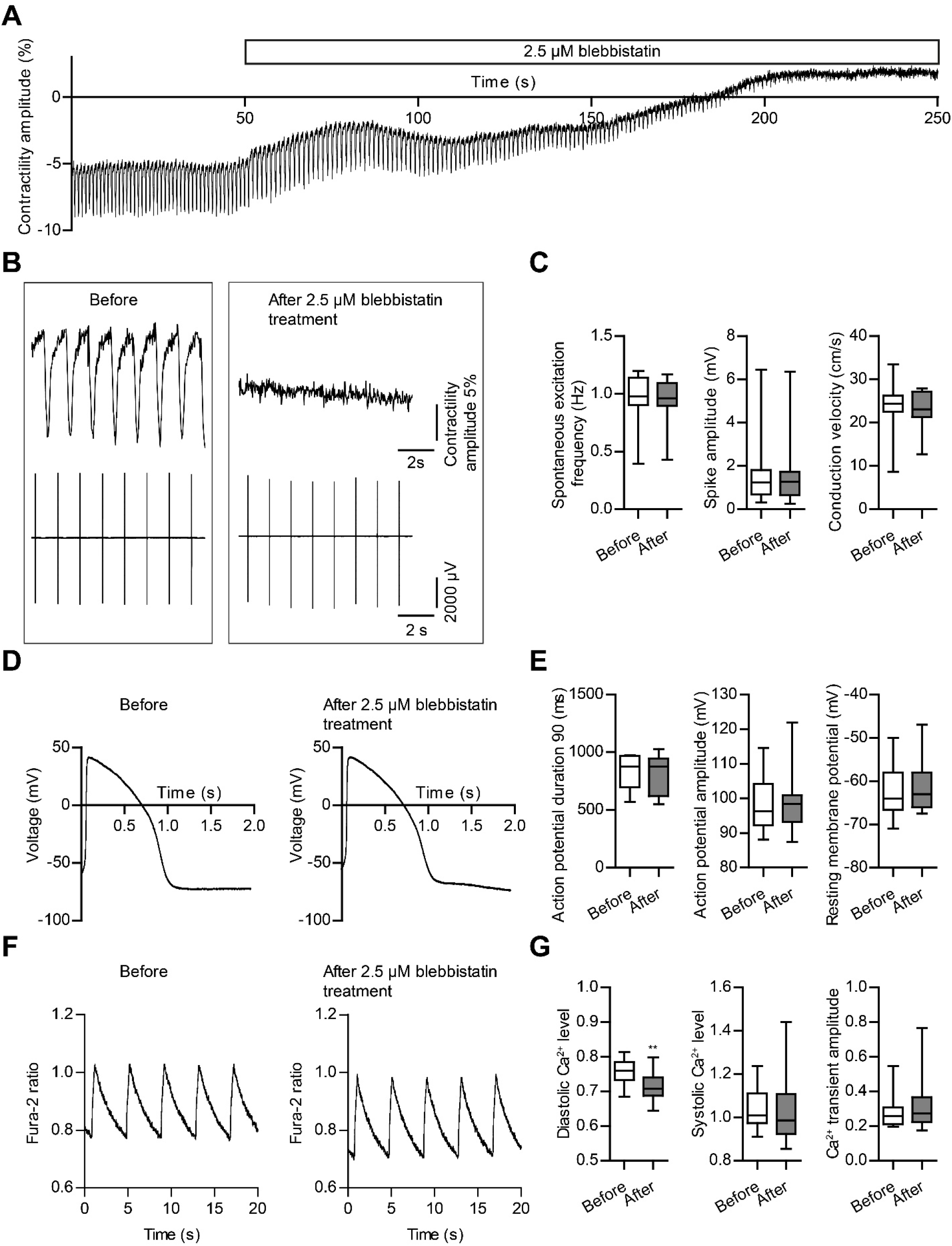
Blebbistatin uncouples the excitation and contraction of iPSC-CMs. (**A**) The representative contractility trace demonstrated the iPSC-CM monolayer culture stopped beating 3–5 minutes after 2.5 μM BB treatment. (**B**) Original contractility and FP traces in the same iPSC-CM culture before and after 2.5 μM BB treatment. (**C**) Statistical analyses of FP metrics: spontaneous excitation frequency (**left**), spike amplitude (**middle**), and conduction velocity (**right**). n = 14 cultures from 5 differentiation experiments for spontaneous excitation frequency and spike amplitude. n = 7 cultures from 3 differentiation experiments for conduction velocity. (**D**) Representative 0.5 Hz paced AP before and after 2.5 μM BB treatment for 10 minutes. (**E**) Statistical analyses of AP metrics: APD_90_ (**left**), AP amplitude (**middle**), and resting membrane potential (**right**). n = 11 cells from 5 differentiation experiments for all the three AP parameters. (**F**) Representative 0.25 Hz paced calcium transients before and after 2.5 μM BB treatment for 5 min. (**G**) Statistical analyses of calcium transient metrics: diastolic Ca^2+^ level (**left**), and systolic Ca^2+^ level (**middle**) and Ca^2+^ transient amplitude (**right**). n = 18 from 4 differentiation experiments for all three calcium transient parameters. Boxplots were used to visualize the spreads of the data. p < 0.05 was considered as significant under the paired Student’s *t*-test (**p < 0.001).

We then investigated the effects of 2.5 μM BB treatment on AP morphologies of iPSC-CMs by using MPC when the cells were paced at 0.5 Hz. We did not observe any differences in APD90, APA, and RMP before and after the treatment with 2.5 μM BB for 10 min (Figure 2D and E). Similar results showing no effect of treatment with 10 μM BB for 5 min on APD were reported in isolated mouse CMs [19]. Furthermore, superfusion of an explanted zebrafish embryonic heart with BB (1, 5 and 10 μM) was reported to have no effects on AP morphology and AP parameters including APD, RMP and maximum upstroke velocity in both atrial and ventricular CMs [30]. Superfusion with 10 μM BB for 60 min also did not induce any changes in AP morphologies paced at 2.5 Hz as registered by microelectrodes from preparations of rabbit atria and ventricles [20]. However, another study showed that the 60-min perfusion with BB (5 μM) significantly prolonged optically recorded APs and corrected QT interval on ECG of the rabbit heart [21].

### Blebbistatin does not alter the magnitude of intracellular calcium transient

To study whether 2.5 μM BB application affects intracellular calcium transient, human iPSC-CMs were labelled with the ratio-metric fluorescent calcium indicator Fura-2 (Figure 2F, G). Our data showed that BB has no effect on Ca^2+^ transient morphology as well as parameters such as amplitude, systolic Ca^2+^ level (Figure 2F, G), or decay rate (before: 1.03 ± 0.015; after: 1.04 ± 0.014; p = 0.576). After 5 min, 2.5 μM BB application led to the reduction of diastolic Ca^2+^ level from 0.76 ± 0.009 to 0.71 ± 0.009 (n = 18 from 4 differentiations, paired, p < 0.001) in iPSC-CMs paced at 0.25 Hz. Similar results were observed in rat cardiomyocytes loaded with the ratio-metric fluorescent calcium indicator Indo-1 [31]. Whereas Ca^2+^ transient amplitude, and the decay were not affected by 1 h application of 0.5 μM BB, diastolic Ca^2+^ level revealed a slight reduction after BB application [31]. Notably, BB application resulted in significant elevations of diastolic fluorescence level in rat cardiomyocytes labelled with the non-ratiometric fluorescent calcium indicators Fluo-4 [31] or Fluo-5F [20] whereas BB did not affect the intracellular Ca^2+^ transient amplitude as assessed by either Fluo-4 or Fluo-5F. Since both Fluo-4 and Fluo-5F are not ratio-metric, therefore, the increase in diastolic fluorescence must be interpreted with caution. Given differences observed by applications of different ratio-metric or non-ratiometric fluorescent indicators, we can speculate that light-sensitive BB tangles with different dyes.

### Blebbistatin protects the function of the sodium channel from calcium paradox

To evaluate whether the relaxation morphologies and state of iPSC-CMs achieved by BB supplement during the whole experimental process (dissociation, straining, storage, and APC recording) make significant differences for current recording, we first recorded *I*_*Na*_ by using physiological external solution containing 140 mM [Na^+^]_o_ (Figure 3A). In the control group without BB, the peak *I*_*Na*_ (−183.4 ± 22.5 pA/pF) appeared at −15 mV while in the group with BB, the peak *I*_*Na*_ of −307.2 ± 35.1 pA/pF (n = 51) was found at −25 mV (Figure 3B). Similarly, the steady-state activation curve was significantly shifted in a direction with more negative membrane potentials in iPSC-CMs with BB buffering compared with iPSC-CMs without BB buffering (Supplementary Figure 2A). V_1/2_ was −36.5 ± 1.2 mV for iPSC-CMs with BB buffering (n = 47), −31.9 ± 1.8 mV for iPSC-CMs without BB buffering (n = 28, P < 0.05). Steady-state inactivation of sodium channels did not differ between the two groups (Supplementary Figure 2B). Unlike the control group without BB, in the group with 2.5 μM BB supplement, the majority of *I*_*Na*_ activation stages lost voltage control (Figure 3A), a sign of too high extracellular Na^+^ concentration for patch-clamp. We even have four ‘out-of-gain’ recordings using the APC gain setting (0.5 mV/pA for a maximum of 20 nA), as we previously observed with MPC using external solution containing 140 mM [Na^+^]_o_, indicating that *I*_*Na*_ recorded in iPSC-CMs with BB treatment were too large at 140 mM [Na^+^]_o_ for the amplifier and voltage clamp was lost. Taken together, these findings suggest that BB supplement protects the sodium channel from Ca^2+^-paradox injury leading to the increase in sodium current density. Therefore, in the following study, we reduced the extracellular Na^+^ concentration and found that none of iPSC-CMs with BB treatment showed ‘out-of-gain’ *I*_*Na*_ under 50 mM [Na^+^]_o_ (Figure 4A-B).

**Figure 3.**
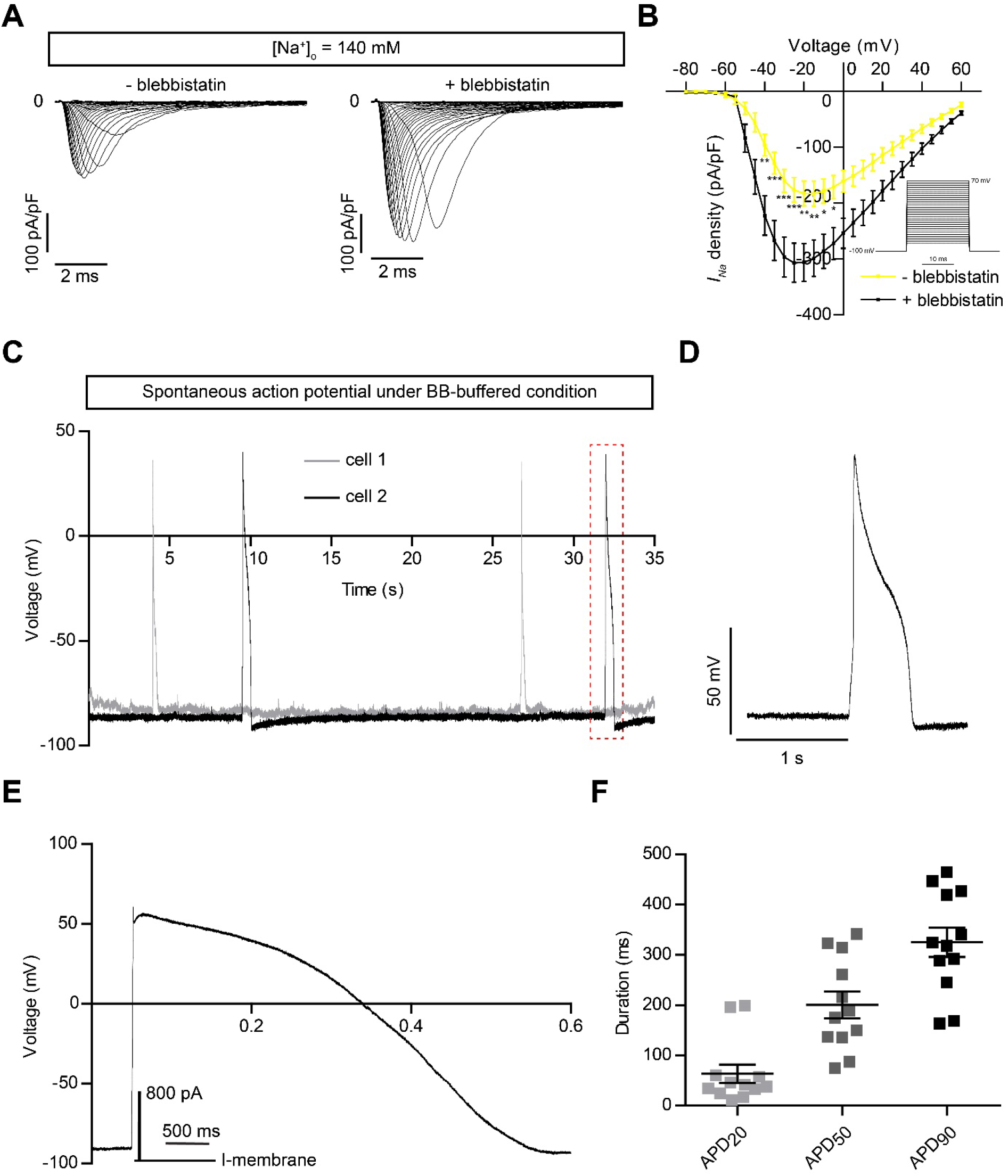
*I*_*Na*_ (140 mM [Na^+^]_o_) recording with/without BB buffering, and AP recording under BB-buffered condition. (**A**) The representative *I*_*Na*_ traces in 140 mM [Na^+^]_o_ buffered with 2.5 μM BB in the whole procedures (including dissociation, filtration, storage, and recording), or without BB. (**B**) Statistical analysis of *I*_*Na*_ in 140 mM [Na^+^]_o_. - blebbistatin: n = 29 cells from 6 differentiation experiments; + blebbistatin: n = 51 cells from 9 differentiations. The stimulation protocol is shown as an inset. (**C**) Dynamic *I*_*K1*_-compensated spontaneous AP traces under BB-buffered condition. (**D**) Enlargement of the spontaneous AP trace of cell 2 in panel C indicated with a red dashed rectangle. (**E**) Original trace of dynamic *I*_*K1*_-compensated, paced AP under BB-buffered condition. Inset: current-clamp protocol for paced AP. (**F**) Scatter plot showing dynamic *I*_*K1*_-compensated paced APD_20_, APD_50_, and APD_90_ under BB-buffered condition (n = 12 cells from 3 differentiations). Data are represented as mean ± SEM. *p < 0.05; **p < 0.01; ***p < 0.001 by two-way ANOVA with Bonferroni post-test.

**Figure 4.**
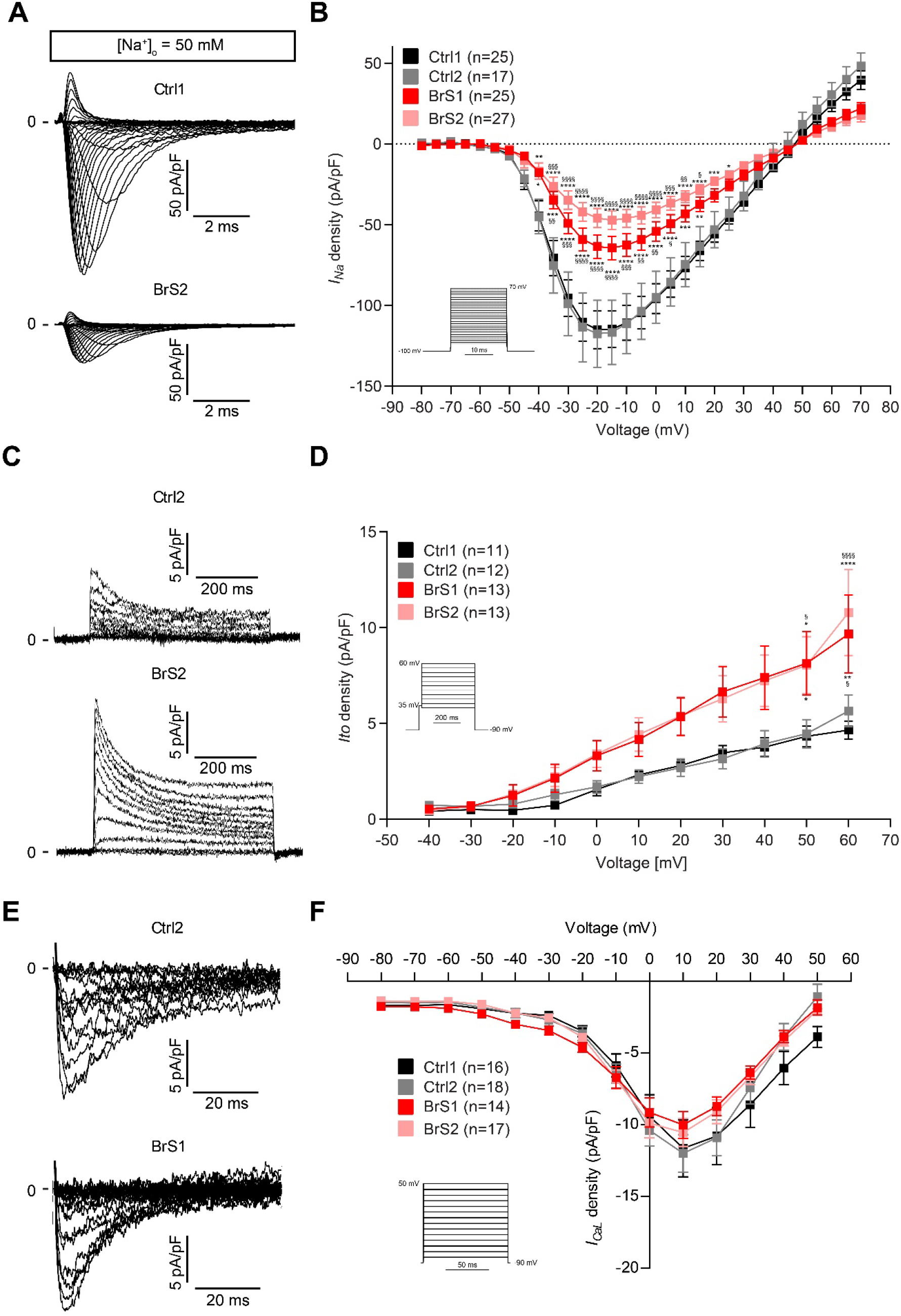
*I*_*Na*_ (50 mM [Na^+^]_o_), *I*_*to*_ and *I*_*CaL*_ recordings for Ctrl- and BrS-CMs buffered with 2.5 μM BB in the whole APC procedure. (**A**) Shown are *I*_*Na*_ recordings (50 mM [Na^+^]_o_) in Ctrl1- and BrS2-CMs. (**B**) The *I-V* curve of *I*_*Na*_ for Ctrl1-, Ctrl2-, BrS1- and BrS2-CMs. Ctrl1: n = 25 cells from 4 differentiation experiments; Ctrl2: n = 17 cells from 3 differentiations; BrS1: n = 25 cells from 7 differentiations; BrS2: n = 27 cells from 7 differentiations. (**C**) Representative traces of *I*_*to*_ in Ctrl2- and BrS2-CMs. (**D**) *I*_*to*_ density plot in Ctrl1-, Ctrl2-, BrS1, and BrS2-CMs. Ctrl1: n = 11 cells from 4 differentiations; Ctrl2: n = 12 cells from 6 differentiations; BrS1: n = 13 cells from 6 differentiations; BrS2: n = 13 cells from 5 differentiations. (**E**) Shown are *I*_*CaL*_ recording in Ctrl2- and BrS1-CMs. (**F**) The *I-V* curve of *I*_*CaL*_ for Ctrl1-, Ctrl2-, BrS1- and BrS2-CMs. Ctrl1: n = 16 cells from 5 differentiations; Ctrl2: n = 18 cells from 7 differentiations; BrS1: n = 14 cells from 4 differentiations; BrS2: n = 17 cells from 3 differentiations. The stimulation protocols are shown as insets. Data are represented as mean ± SEM. *p < 0.05; **p < 0.01; ***p < 0.001; ****p < 0.0001 BrS *vs*. Ctrl1 by using two-way ANOVA with Bonferroni post-test. ^§^p < 0.05; ^§§^p < 0.01; ^§§§^p < 0.001; ^§§§§^p < 0.0001 BrS *vs*. Ctrl2 by using two-way ANOVA with Bonferroni post-test.

### The use of blebbistatin enables the recording of spontaneous I_K1_-compensated APs with APC

A big problem for electrophysiological studies using human iPSC-CMs with APC is that it is almost impossible to record spontaneous APs, even after an injection of virtual, simulated *I*_*k1*_ into the cells [9], because of fresh enzymatic digestion and Ca^2+^-paradox injury. Under BB-buffered condition, we were surprised to obtain some CMs exhibiting spontaneous APs with typical morphology after an injection of virtual, simulated *I*_*k1*_ into the cells (Figure 3C and D). Compared to our previously published APD_50_ and APD_90_ data of dynamic *I*_*K1*_-supplemented, paced APs in iPSC-CMs without BB buffering [9], APD_50_ and APD_90_ were significantly longer (APD_50_: 200.4 ± 26.4 ms *vs*. 108.8 ± 22.7 ms, p = 0.017, *t*-test; APD_90_: 324.8 ± 29.3 ms *vs*. 193.7 ± 28.9 ms, p = 0.005, *t*-test) in iPSC-CMs under BB-buffered condition (Figure 3E and F). In cardiomyocytes, Ca^2+^ influx via L-type calcium channels plays a multi-functional role in controlling APD and regulating muscle contraction [32]. Considering that BB itself does not affect APD of iPSC-CMs (Figure 2D and E), it is possible that BB protects the function of not only the sodium channel but also other ion channels (e.g. L-type calcium channel and potassium channels) from Ca^2+^-paradox injury in single iPSC-CMs during the APC process. Previous studies demonstrated that calcium paradox injury was also mediated through transient receptor potential canonical (TRPC) channels in mouse ventricular myocytes [13] and sevoflurane can protect cardiomyocytes from calcium paradox-mediated calcium overload by blocking the activation of TRPC channels [33]. It would be interesting to use TRPC blockers (like sevoflurane) to intervene the calcium paradox in future studies. Taken together, these findings indicate that hypercontracted iPSC-CMs are not suitable for electrophysiological research using APC because the calcium paradox can have catastrophic consequences without any intervention [11, 14, 15]. Therefore, electrophysiological studies using human iPSC-CMs with APC require the use of excitation-contraction coupling uncoupler during single cell preparation and for current and AP recordings.

### Blebbistatin facilitates disease modelling for Brugada syndrome using iPSC-CMs with APC

To study whether human iPSC-CMs under BB-buffered condition can be used with APC for disease modelling, we repeated *I*_*Na*_ recording under 50 mM [Na^+^]_o_ using APC in previously published BrS disease models [34]. In the presence of 2.5 μM BB, single iPSC-CMs were obtained from two healthy donors (Ctrl1 = iBP76.3 and Ctrl2 = iWT.D2.1) and two BrS patients (BrS1 and BrS2 harboring the same *SCN5A* p.S1812X mutation. The APC system Patchliner Quattro used in this study allows the recordings from 4 cells simultaneously. To reduce the variabilities among different cell lines and experiments, we performed at least 3 differentiation experiments for each donor/patient and carried out recordings from 4 or 8 cells per experiment. As shown in Figure 4A, the *I*_*Na*_ density in BrS-CMs was dramatically lower compared to that in Ctrl-CMs. Under the testing potentials ranging from −40 mV to 15 mV, *I*_*Na*_ densities in both BrS1- and BrS2-CMs were significantly smaller than in Ctrl1- and Ctrl2-CMs (Figure 4B). The peak *I*_*Na*_ appeared at −20 mV in Ctrl1- and Ctrl2-CMs showed as −114.9 ± 11.7 and −117.5 ± 20.7 pA/pF, and in BrS1- and BrS2-CMs presented as −63.5 ± 7.3 and −45.9 ± 6.0 pA/pF, respectively. These results generated by using APC were consistent with our previous results acquired with MPC, which also revealed a more than 50% reduction of *I*_*Na*_ density in BrS-CMs [34].

The peak *I*_*to*_ at +60 mV in BrS1-CMs (9.7 ± 2 pA/pF) and BrS2-CMs (10.8 ± 2.2 pA/pF) were significantly bigger than those in Ctrl1-CMs (4.7 ± 0.5 pA/pF) and Ctrl2-CMs (5.7 ± 0.8 pA/pF) (Figure 4C and D). The *I*_*to*_ recorded with APC are in line with our previous publication of MPC results: the *I*_*to*_ at +60 mV in BrS1-CMs and BrS2-CMs were 2.4 and 1.9 times bigger than those in Ctrl-CMs [34]. Furthermore, consistent with our MPC data, we did not observe any *I*_*CaL*_ density differences between Ctrl-CMs and BrS-CMs (Figure 4 E, F).

In summary, we first report the optimization of single iPSC-CM preparation for APC recording in this study. With the supplement of BB to the whole procedures (dissociation, filtration, storage, and recording), we could make significant improvement, not only in obtaining relaxed iPSC-CMs, but also in *I*_*Na*_ and AP recordings. Furthermore, APC faithfully recapitulates the single-cell electrophysiological phenotypes of iPSC-CMs derived from BrS patients, as detected with MPC. Our study suggests that APC is capable of replacing MPC in the modeling of cardiac channelopathies using human iPSC-CMs by providing high-quality data with higher throughput. Although the high cost of APC consumables (e.g. chips, plates, high number of iPSC-CMs needed) still limits its wide application in channelopathy research and drug screening, the development of APC technology is continuing to meet the needs across drug discovery in pharmaceutical, biotech and academic research laboratories. For example, the new SyncroPatch 384 APC system enables parallel data acquisition of 384 cells and is suitable for high-throughput research.

## Author Contributions

WL and KG conceived the study and designed experiments. WL, XL, and YU performed experiments and acquired data. WL, XL, and KG analyzed and interpreted the data. WL and KG wrote the manuscript.

## Acknowledgments

We thank Konstanze Fischer, Jessie Pöche, and Judith Müller for excellent technical assistance. The authors would like to express great appreciation to Free State of Saxony and the European Union EFRE for project funding (SAB projects “PhänoKard” and “PhenoCor” to K.G.).

