## Supplemental Figure 1 for "Blebbistatin protects iPSC-CMs from hypercontraction and facilitates automated patch-clamp based electrophysiological study"

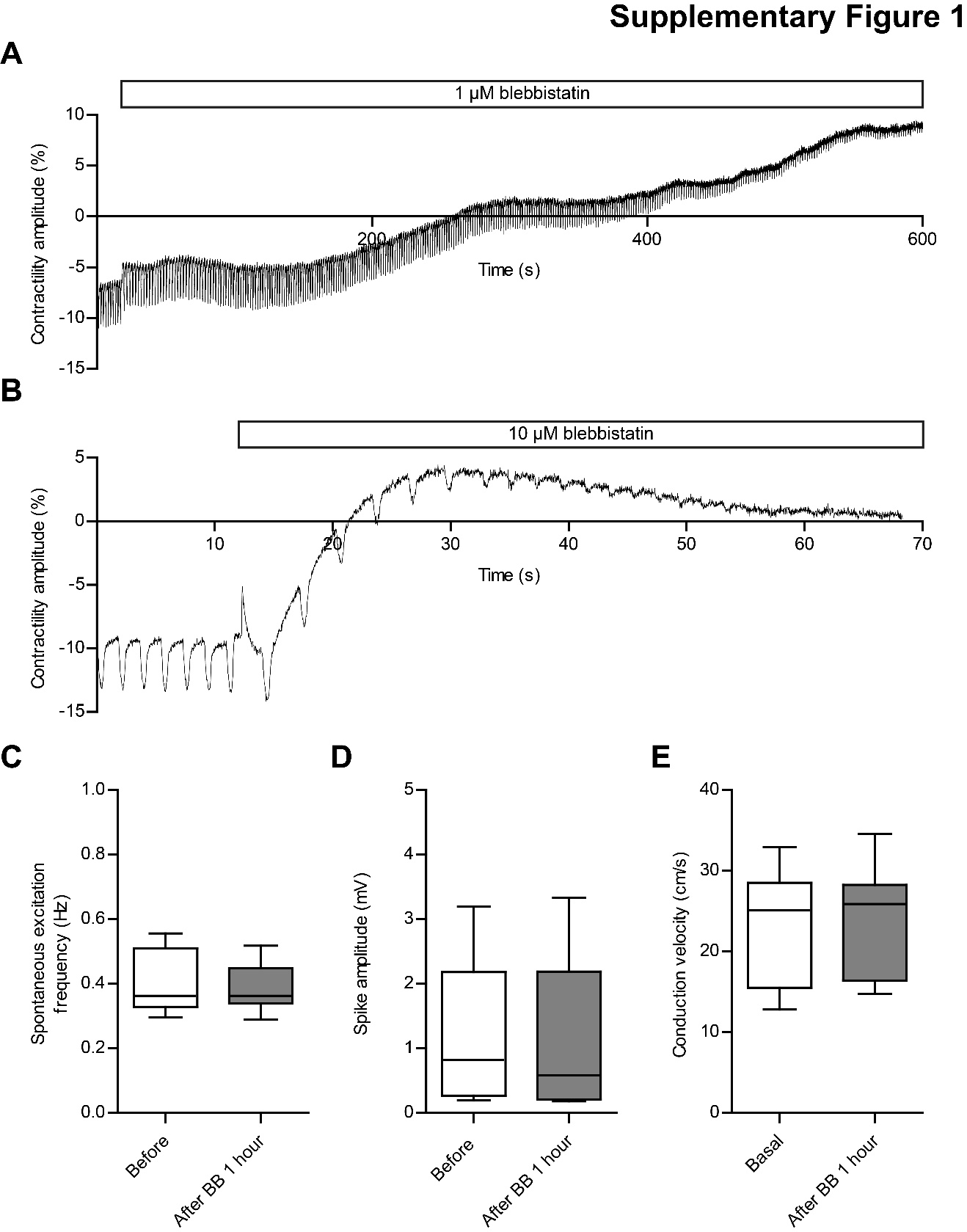

**Supplementary Figure 1. Blebbistatin shows no effects on FP signals of iPSC-CMs.** (**A-B**) Original contractility traces shown the inhibiting effects of 1 µM (**A**) and 10 µM (**B**) blebbistatin. **(C-E)** Statistical analyses of FP metrics: spontaneous excitation frequency (**C**), spike amplitude (**D**), and conduction velocity (**E**) after 1 hour treatment with 2.5 µM BB. n = 18 cultures from 3 differentiation experiments for spontaneous excitation frequency and spike amplitude. n = 8 cultures from 2 differentiation experiments for conduction velocity. Boxplots were used to visualize the spreads of the data. p < 0.05 was considered as significant under the paired Student’s *t*-test.

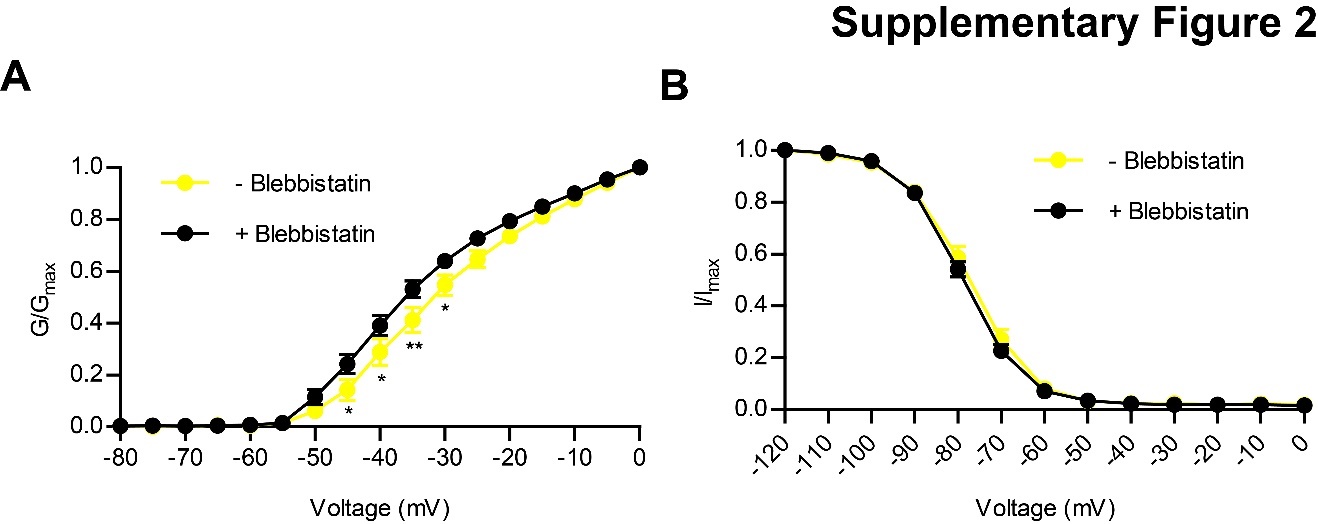

**Supplementary Figure 2. *I_Na_* Steady-state activation and inactivation of** ***I_Na_* (140 mM [Na^+^]_o_) recording with/without blebbistatin buffering.** (**A**) Steady-state activation. - blebbistatin: n = 28 cells from 6 differentiation experiments; + blebbistatin: n = 47 cells from 9 differentiations. (**B**) Steady-state inactivation. - blebbistatin: n = 24 cells from 5 differentiation experiments; + blebbistatin: n = 43 cells from 9 differentiations. Two-way ANOVA with Bonferroni post-hoc test was used for statistical analysis (* p < 0.05, ** p < 0.01).

| **Substance**  **(mM)** | **MPC solutions** | | **APC solutions** | | | | | | | | |
| --- | --- | --- | --- | --- | --- | --- | --- | --- | --- | --- | --- |
|  | APs | | External solution | Seal Enhancer | *I_to_* and APs | | *I_Na_* | | | *I_CaL_* | |
|  | Bath (Tyrode) | Pipette | For cell catching | For seal | Bath | Pipette | Bath under 50 mM [Na^+^]_o_ | Bath under 140 mM [Na^+^]_o_ | Pipette | Bath | Pipette |
| NaCl | 138 |  | 140 | 130 | 140 | 10 | 50 | 140 | 10 |  | 10 |
| Na_2_-phosphocreatine |  | 5 |  |  |  |  |  |  |  |  |  |
| TEA-Cl |  |  |  |  |  |  | 90 |  |  | 150 |  |
| KCl | 4 | 60 | 4 | 4 | 4 | 10 |  | 4 |  |  |  |
| K_D_-gluconate |  | 100 |  |  |  |  |  |  |  |  |  |
| CaCl_2_ | 1.8 |  | 2 | 10 | 2 |  | 2 | 2 |  | 2 |  |
| MgCl_2_ | 1 |  | 1 | 1 | 1 |  | 1 | 1 |  | 1 |  |
| Mg-ATP (freshly) |  | 4 |  |  |  | 1 |  |  | 1 |  | 1 |
| NaGTP  (freshly) |  | 0.3 |  |  |  |  |  |  |  |  |  |
| CsCl |  |  |  |  |  |  | 4 |  | 30 |  | 30 |
| KF |  |  |  |  |  | 110 |  |  |  |  |  |
| CsF |  |  |  |  |  |  |  |  | 90 |  | 90 |
| Glucose | 10 |  | 5 | 5 | 5 |  | 5 | 5 |  | 5 |  |
| HEPES | 10 | 10 | 10 | 10 | 10 | 10 | 10 | 10 | 10 | 10 | 10 |
| EGTA |  |  |  |  |  | 10 |  |  | 10 |  | 10 |
| NaH_2_PO_4_ | 0.33 |  |  |  |  |  |  |  |  |  |  |
| Nifedipine |  |  |  |  |  |  | 0.01 | 0.01 |  |  |  |
| CdCl_2_ |  |  |  |  | 0.5 |  |  |  |  |  |  |
| pH adjustment | 7.3  with  NaOH | 7.2  with  KOH | 7.4  with  NaOH | 7.4  with  NaOH | 7.4  with  NaOH | 7.2  with  CsOH | 7.4  with  NaOH | 7.4  with  NaOH | 7.2  with  CsOH | 7.3  with  CsOH | 7.2  with  CsOH |

**Supplementary Table 1. The formula of extracellular and intracellular solutions for manual and automated patch-clamp.**
